# Structure and function of a bacterial gap junction analog

**DOI:** 10.1101/462465

**Authors:** Gregor L. Weiss, Ann-Katrin Kieninger, Iris Maldener, Karl Forchhammer, Martin Pilhofer

## Abstract

Multicellular lifestyle requires cell-cell connections. In multicellular cyanobacteria, septal junctions enable molecular exchange between sister cells and are required for cellular differentiation. The structure of septal junctions is poorly understood and it is unknown whether they regulate intercellular communication.

Here we resolved the *in situ* architecture of septal junctions by electron cryotomography of cryo-focused ion beam-milled cyanobacteria. Septal junctions consisted of a tube traversing the septal peptidoglycan. Each tube end comprised a plug that was covered by a cytoplasmic cap. Fluorescence recovery after photobleaching showed that intercellular communication was blocked upon stress. This gating was accompanied by a conformational change of the septal junctions, mediated by the proteins FraC/D.

We provide the mechanistic framework for a cell junction that predates eukaryotic gap junctions by a billion years. The conservation of a gated dynamic mechanism across different domains of life emphasizes the importance of controlling molecular exchange, e.g. upon injury.

## Introduction

The evolution of multicellular organisms allowed to divide specialized tasks among sister cells and led to the invention of structures mediating intercellular molecular exchange (Brunet and King, 2017). Metazoan cells communicate via gap junctions, which are multimeric protein complexes that form two hemi-channels and can control molecular exchange by a dynamic conformational change (Hervé and Derangeon, 2013; Unwin and Zampighi, 1980). In plants, plasmodesmata generate continuity between the cytoplasm of neighboring cells, however, they are mainly composed of membranes, their structure is highly heterogeneous, and closing is possible by polysaccharide (callose) deposition on a time scale of only hours to days (Oparka et al., 1999; Sager and Lee, 2014).

Filamentous cyanobacteria are true multicellular organisms. Under nitrogen limiting conditions, strains of the order *Nostocales* differentiate N_2_-fixing heterocysts in a semiregular pattern along the filament, which supply the neighboring vegetative cells with nitrogen-fixation products in form of glutamine and dipeptide β-aspartyl-arginine (Burnat et al., 2014; Thomas et al., 1977). Vegetative cells, in turn, fix CO_2_ via oxygenic photosynthesis and provide heterocysts with sucrose as a carbon and energy source (Cumino et al., 2007; Jüttner, 1983). In addition to metabolites, signaling molecules need to be exchanged to establish the correct pattern of differentiated cells along the filament.

Transport between neighboring cells occurs by diffusion through the septa, as shown by tracer molecules like fluorescent calcein with a weight of ~620 Da (Flores et al., 2016; Mullineaux et al., 2008; Nieves-Morión et al., 2017). The exchange of molecules through the septal peptidoglycan (PG) cell wall requires an array of 80 to 150 nanopores, generated by cell wall-lytic amidases in the model organisms *Nostoc punctiforme* and *Anabaena* sp. PCC 7120 (here *Anabaena*) (Bornikoel et al., 2017; Lehner et al., 2013). It has been speculated that the nanopores, together with any of the septal proteins SepJ, FraC and FraD, form so-called ‘septal junctions’ (SJs). SJs might establish a direct connection between the cytoplasm of neighboring cells, spanning the entire periplasmic space (Flores et al., 2016; Wilk et al., 2011). Even though cyanobacterial SJs represent an ancient type of cell junction, their structure and composition, as well as whether they can control cell-cell communication is unknown.

## Results and Discussion

### *In situ* architecture of septal junctions reveals tube, plug and cap modules

Here, we imaged *Anabaena* cells by electron cryotomography (ECT) to reveal the architecture of SJs *in situ* and in a near-native state. To obtain a sample that was thin enough for ECT imaging, we plunge-froze cells on electron microscopy (EM) grids and prepared lamellae using cryo-focused ion beam (FIB) milling (Figure S1) (Marko et al., 2007; Medeiros et al., 2018; Rigort et al., 2010; Schaffer et al., 2017). Tomograms of septa between vegetative cells revealed numerous putative SJs that appeared as tubular structures traversing the septum (Figure 1a/b; Movie S1). In a 200 nm thick lamella, an average of 9.8 SJs were clearly visible (n=21 tomograms), consistent with the reported number of ~80 nanopores in a septum (Bornikoel et al., 2017). Structures resembling SJs were never observed in the lateral cell wall. The cross-sectional density plot of a SJ suggests that a tube structure is inserted into the septal PG (rather than the PG channel being empty) and the tube lumen density was relatively low compared to the PG (Figure 2). Depending on the thickness of the septum, the length of the tube module varied between 26 and 79 nm (average 37.9 nm +/-7.1 nm, n=208, Figure S3), suggesting a multimeric nature of the tube.

**Figure 1.**
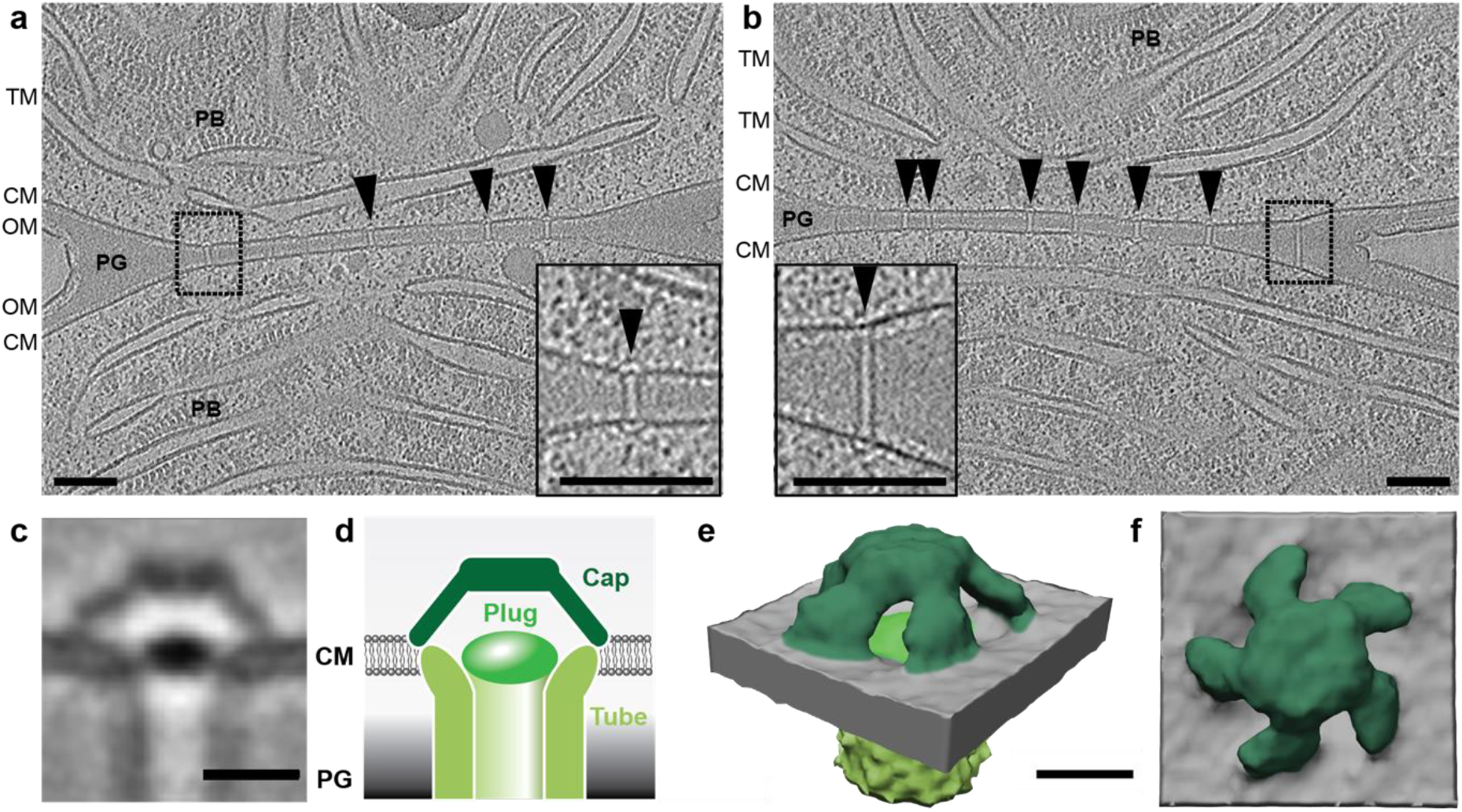
*In situ* architecture of septal junctions reveals tube, plug and cap modules. **a, b:** Cryotomograms (magnified views in boxes) of a FIB-milled *Anabaena* filament. The two different slices at different Z-heights show the septum between adjacent vegetative cells. Multiple SJs are seen crossing the septum (arrowheads), which are precisely adjusted to the septum thickness. CM, cytoplasmic membrane; OM, outer membrane; PB, phycobilisomes; PG, septal peptidoglycan; TM, thylakoid membranes. Bars, 100 nm. Shown are projections of 13.5 nm thick slices. **c-f**: Subtomogram averaging of SJ ends revealed three structural modules: tube, cap and plug. Shown is a 0.68 nm-thick tomographic slice through the average (c), a schematic representation of SJ modules (d; modules segmented in different shades of green), and oblique (e) and top (f) views of a surface representation (modules were segmented to match colors in d). The cap consisted of a ceiling that was held by five arches. Bars, 10 nm.

**Figure 2.**
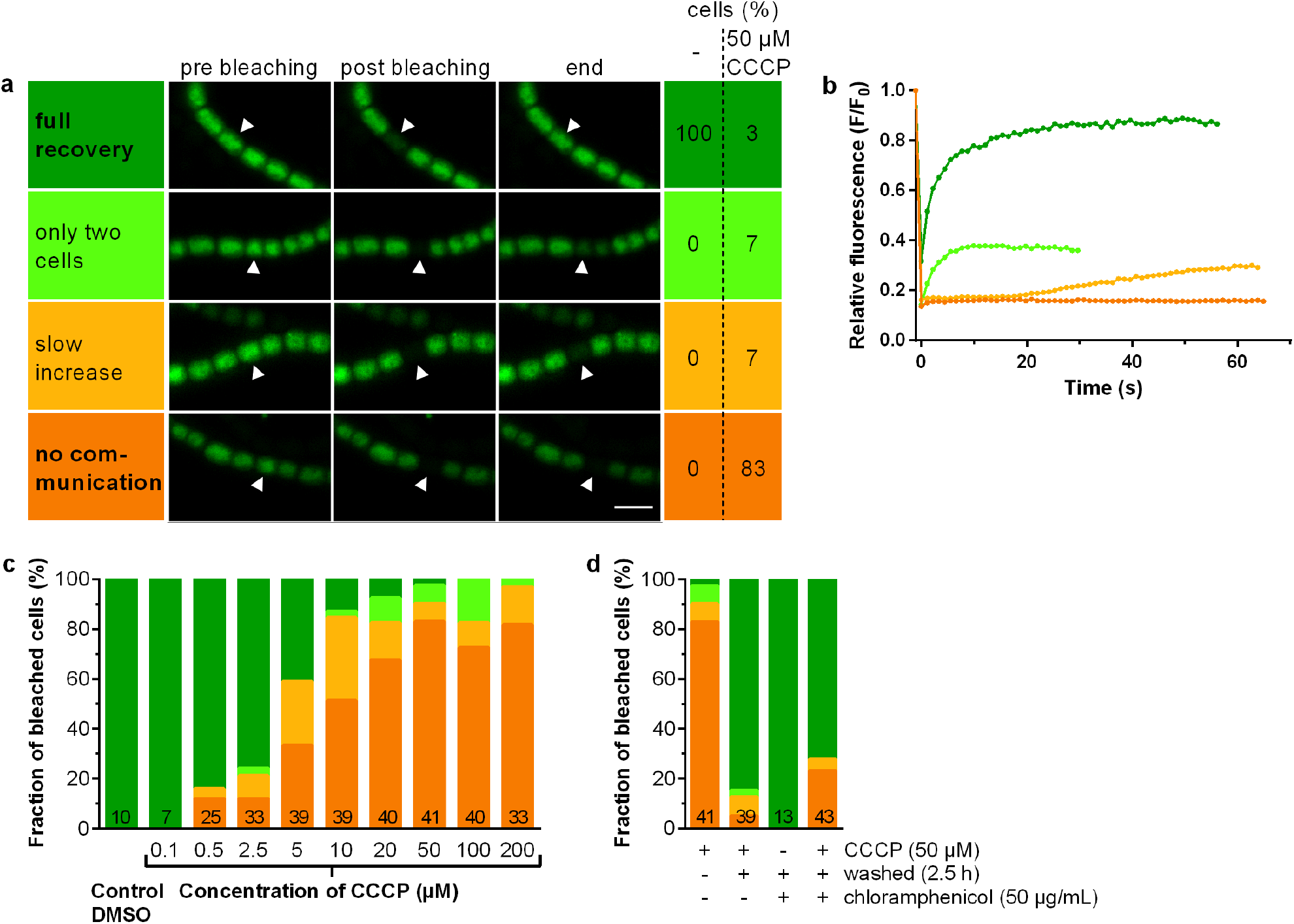
Intercellular communication ceases upon ionophore treatment in a reversible manner. **a:** Shown is FRAP analysis of cells that were stained with fluorescent calcein. In the control experiment (DMSO), all cells showed full recovery of fluorescence after bleaching. When the culture was treated with the ionophore CCCP (50 μM), the bleached cells showed four different types of FRAP responses: ‘no communication’ (no recovery), ‘slow increase’ (delayed recovery to <50% of original fluorescence), ‘only two cells’ (exchange of calcein only with one neighboring cell), and ‘full recovery’. For each FRAP response, representative images are shown at three time points (5 s before bleaching, ~0.5 s after bleaching, 30-60 s after bleaching). Arrowheads point to the bleached cells. *Anabaena* was apparently able to control communication upon disruption of the membrane potential, since the majority of CCCP-treated cells showed ‘no communication. Bar 5 μm. **b**: Shown are fluorescence recovery curves corresponding to the four FRAP responses that were observed in ‘a’ (color scheme identical to ‘a’). Time point t=0 shows the analyzed cell directly after bleaching. **c**: The quantification of FRAP responses (color scheme identical to ‘a’) indicates that the effect of CCCP on cell-cell communication was concentration-dependent for CCCP concentrations between 0.5-50 μM. In the control experiment, cells were treated with 0.002% DMSO. Numbers within the bars indicate the number of analyzed cells *n* from different filaments and represent cumulated results from at least 2 independent cultures (except for 0.1 μM CCCP). Results of independent cultures are shown in Figure S6a. **d**: CCCP-mediated control of cell-cell communication was reversible, since communication was observed after incubation in fresh medium lacking CCCP (color scheme identical to ‘a’). Regaining cell-cell communication was independent of *de novo* protein synthesis, shown by the experiment that was conducted in the presence of chloramphenicol (inhibiting protein synthesis). ‘+’ and ‘-’ indicate the presence and absence of CCCP, washing in fresh medium, and chloramphenicol. Numbers in bars indicate number of analyzed cells *n*. Shown are cumulated results from at least two independent cultures. Results of independent cultures are shown in Figure S6b.

In addition to the tube module, the tomograms revealed a cytoplasmic cap-like structure, as well as a plug-like density in the cytoplasmic membrane (CM; Figure 1a/b). Both ends of each SJ comprised cap and plug modules, without any recognizable differences between both ends. To increase contrast and resolution, we performed subtomogram averaging of 446 SJ ends (Figure 1c-f; Figure S4, Movie S2). The average resolved that the 11 nm-wide tube (lumen of 7 nm) made direct contact with the CM. In contrast to the resolved lipid bilayer in the CM, no bilayer-like density was observed in the SJ tube wall. This supports the idea of a proteinaceous building block assembling a multimeric, periplasma-spanning tube and is consistent with earlier reports (Wilk et al., 2011). The plug (7 nm x 2.5 nm) was sitting at the end of the tube at the level of the CM. The cap module was a 5-fold rotational symmetric (Figure S5) dome (8 nm height) covering the tube end. The ceiling had a diameter of 9.5 nm and was held by five arches with lengths of 8.5 nm.

### Intercellular communication ceases upon ionophore treatment in a reversible manner

The structural complexity of the SJ ends led us to speculate that the assembly might allow the control of intercellular molecular diffusion. We therefore monitored the molecular exchange rate upon challenging the membrane potential. Cyanobacterial intercellular communication was studied previously by monitoring the exchange rate of fluorescent calcein by fluorescence recovery after photobleaching (FRAP) (Mullineaux et al., 2008; Nürnberg et al., 2015). Here, before FRAP analysis, we treated cells with carbonyl cyanide 3-chlorophenylhydrazone (CCCP), a protonophore that uncouples the proton gradient across membranes (Hopfer et al., 1968). Upon treatment with 50 μM CCCP, 83% of the analyzed cells ceased to exchange calcein, showing a ‘no communication’ response (Figure 2a/b; Movie S3). This is in contrast to the control experiment, where all DMSO-treated cells displayed ‘full recovery’ of fluorescence (Figure 2a). Further 7% of CCCP-treated cells were assigned to a ‘slow increase’ response, since fluorescent recovery was delayed (started only 20-60 s after bleaching) and reached only <50% of the initial fluorescence (Figure 2a/b). In 7% of the cells, exchange took place only with a single neighboring cell. Only 3% of CCCP-treated cells showed a normal ‘full recovery’ FRAP response (Figure 2a/b). The fraction of non-communicating cells was CCCP concentration-dependent (Figure 2c, Figure S6a), having no effect below 0.5 μM. Concentrations above 50 μM did not further inhibit molecular exchange.

To test whether CCCP inhibited cell-cell communication in a reversible manner, cells were washed after a 50 μM CCCP treatment and incubated in fresh medium for 2.5 h at room temperature. Eighty-five percent of the cells resumed communication, suggesting that the inhibitory control mechanism was indeed reversible (Figure 2d, Figure S6b). Cells were also still viable after CCCP treatment (Figure S7). We then set out to explore whether the re-opening of SJs required the synthesis of new proteins. Hence, cells were treated with CCCP, washed, and incubated for 2.5 h in fresh media supplemented with 50 μg/mL chloramphenicol (inhibiting protein synthesis) before monitoring the FRAP response (Figure 2d, Figure S6b). Since 72% of the tested cells were able to restore communication (showing ‘full recovery’ response), we concluded that the reversibility of communication was based on an opening mechanism of SJs that was independent of *de novo* protein synthesis.

### Ceased intercellular communication after ionophore treatment coincides with a major structural rearrangement of the septal junction cap

To investigate whether a structural change in the macromolecular architecture of SJs was involved in the gating of cell-cell communication, we plunge froze CCCP-treated *Anabaena* cells and acquired tomograms of septal areas. Differences were hardly detectable in individual tomograms (Figure 3a). However, subtomogram averaging revealed a striking conformational change in the cap module, whereas the tube and plug modules remained unchanged (Figure 3b/c/d; Movie S4). Compared to the cap structure in untreated cells (Figure 3e), the individual arches were not anymore detectable and the cap did not reveal any detectable openings (Figure 3b, Figure S8; Movie S5). The structural rearrangement also resulted in a tightening of the cap by 6 nm and the introduction of a small cavity on the ceiling of the cap. It is possible that the closed confirmation of the cap could arise from rotations of the individual arches (Figure S9).

**Figure 3.**
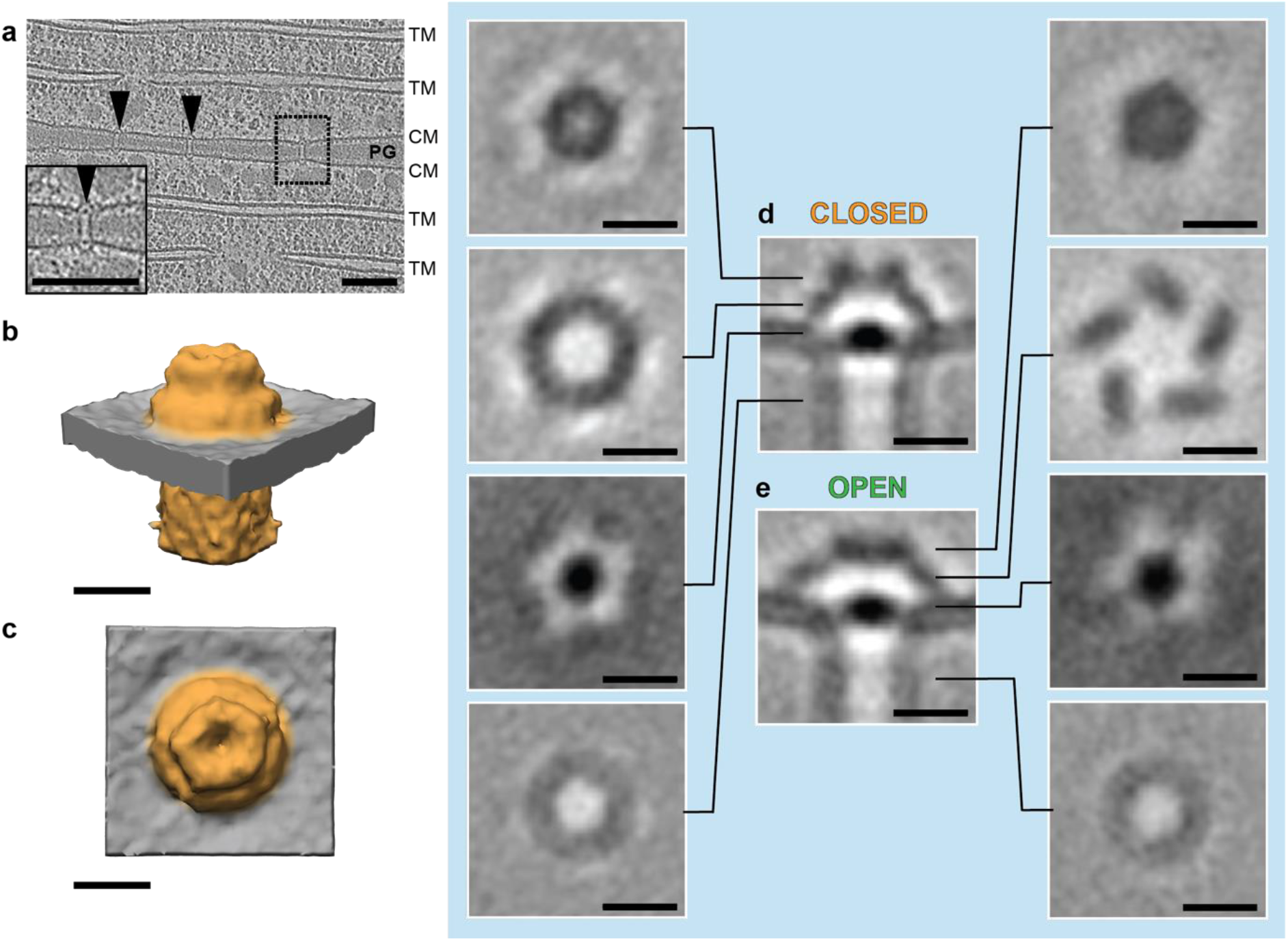
Ceased intercellular communication after ionophore treatment coincides with a major structural rearrangement of the septal junction cap. **a**: Shown is a 13.5 nm-thick slice through a cryotomogram (magnified view in box) of the septal area of a CCCP-treated *Anabaena* filament. SJs are indicated by arrowheads. CM, cytoplasmic membrane; PG, septal peptidoglycan; TM, thylakoid membranes. Bar, 100 nm. **b-e**: Subtomogram averaging of SJs in the CCCP-treated non-communicating ‘closed’ state (b-d) revealed major structural rearrangements in the cap module, compared to the ‘open’ state (e). Shown are surface representations (b/c), and longitudinal and cross-sectional slices (0.68 nm) through the averages (d/e). Sliced positions are indicated in d/e. Bars, 10 nm.

### The cap and plug modules are required to control intercellular communication upon ionophore treatment

Since the conformational switch of SJs from ‘open’ to ‘closed’ coincided with the loss of intercellular molecular diffusion, we analyzed mutants to further explore the involvement of the different modules in controlling communication. AmiC1, FraC and FraD were proposed to play important roles in the formation or as structural components of SJs (Flores et al., 2016). We therefore analyzed respective mutants by ECT (Figure 4a/b/c) and we also tested their ability to control intercellular molecular exchange upon ionophore treatment (Figure 4d). The number of SJs was significantly reduced in all tested mutants (*amiC1* mutant SR477, Δ*fraD* and Δ*fraC-*Δ*fraD*), which is consistent with previous quantifications of nanopore arrays (Bornikoel et al., 2017; Nürnberg et al., 2015). The septa of all mutants were also wider, which was reflected in the increased SJ average length (Figure S10). A subtomogram average of the *amiC1* mutant SR477 (Figure 4a) showed that SJs were in the open state and did not reveal any structural differences compared to the wildtype. When we monitored intercellular molecular exchange by FRAP, we found that only 52% of SR477 cells showed ‘full recovery,’ likely based on the low number of SJs. Upon CCCP treatment, 87% of the analyzed cells showed a ‘no communication’ response (Figure 4d, Figure S6c), suggesting that the low number of SJs could mostly still switch to the closed state. The amidase AmiC1 is therefore unlikely a direct structural component of SJs, consistent with previous assumptions (Bornikoel et al., 2017; 2018; Lehner et al., 2013; Nürnberg et al., 2015).

**Figure 4.**
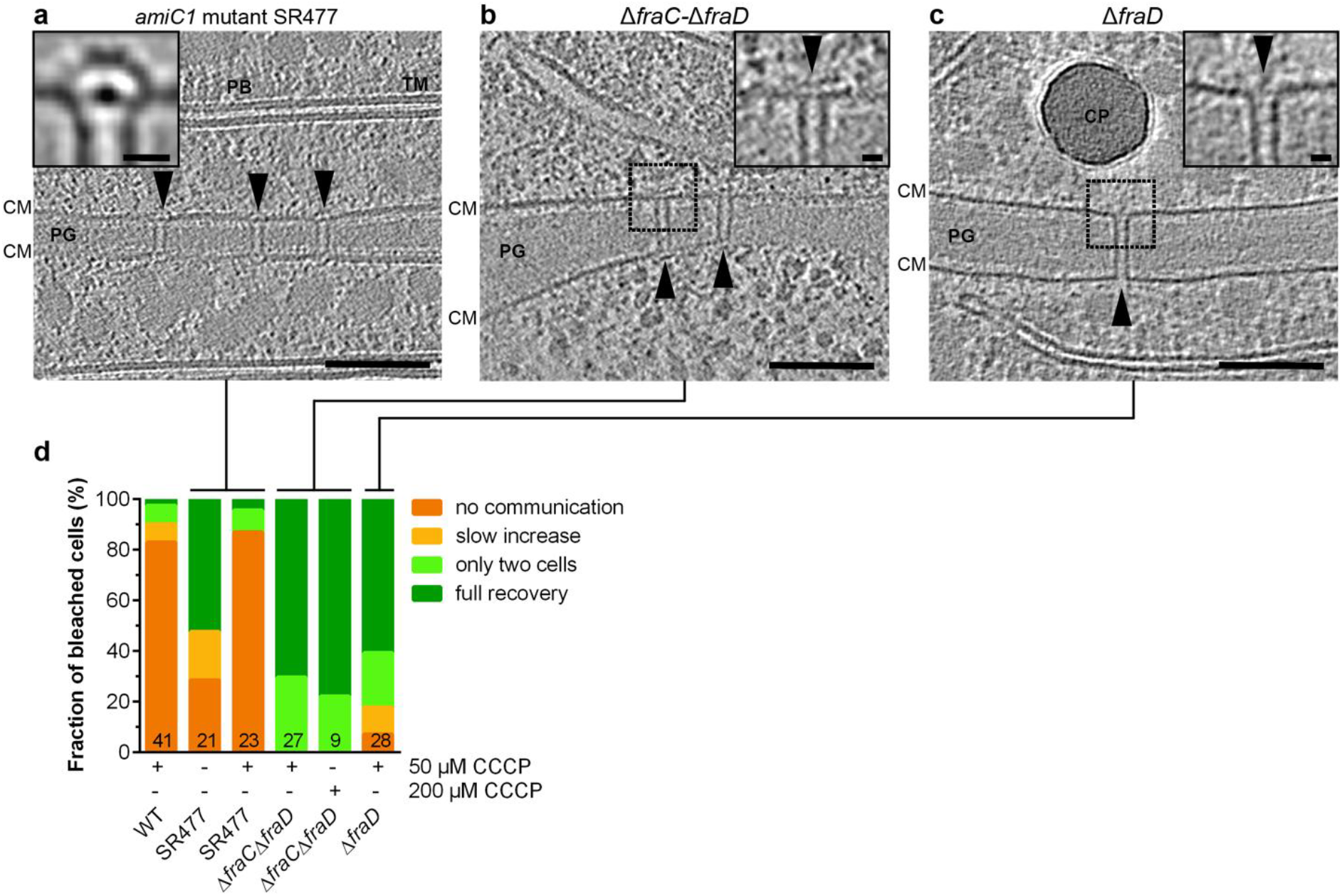
The cap and plug modules are required to control intercellular communication upon ionophore treatment. **a-c:** The cryotomograms (shown are 13.5 nm-thick projections, bars 100 nm) of different *Anabaena* mutants showed that the number of SJs (arrowheads) was significantly reduced. SJs from the *amiC1* mutant SR477 were in the open state and revealed no differences compared to the wildtype (a, inset shows subtomogram average; 0.68 nm thick slice; bar 10 nm). SJs from the Δ*fraC-*Δ*fraD* (b) and Δ*fraD* (c) mutants were missing the cap and plug modules (insets show magnified views, bars 10 nm). CM, cytoplasmic membrane; CP, cyanophycin; PB, phycobilisomes; PG, septal peptidoglycan; TM, thylakoid membranes. **d:** FRAP responses of the *amiC1* mutant SR477 showed that compared to the wildtype, a reduced fraction of cells was able to communicate already in the absence of CCCP (likely based on the lower total number of SJs). However, the open SJs of SR477 were able to close upon CCCP treatment, consistent with the presence of cap/plug in the structure. The Δ*fraD* and Δ*fraC-*Δ*fraD* mutants were unable to close SJs upon CCCP treatment, consistent with the absence of cap/plug modules. Numbers within the bars indicate the number of analyzed cells *n* from different filaments. Results from at least two independent cultures were cumulated (except for Δ*fraC-*Δ*fraD* treated with 200 μM CCCP). Results of independent cultures are shown in Figure S6c.

The SJs of the Δ*fraC-*Δ*fraD* and Δ*fraD* mutants lacked the cap and plug modules (Figure 4b/c). Importantly, none of the analyzed Δ*fraC-*Δ*fraD* cells and only 7% of the Δ*fraD* cells showed a ‘no communication’ response upon CCCP treatment, even at high CCCP concentrations (Figure 4d, Figure S6c). Together, these data suggest that cap and/or plug are required to close SJs and thereby terminate intercellular molecular diffusion. Interestingly, FraD comprises five predicted transmembrane helices and a C-terminal periplasmic segment (Merino-Puerto et al., 2011). Therefore, FraC and FraD could represent a structural SJ element, and/or be required for plug/cap assembly.

Besides our finding that SJs closed upon the disruption of the membrane potential, we also tested intercellular communication in the presence of oxidative stress by treatment with 10 mM H_2_O_2_ for 3 h. Similar to the CCCP treatment, the fraction of cells showing the FRAP response ‘full recovery’ dropped from 100% to 6% (Figure S11). Consistent with the above data, the Δ*fraC-*Δ*fraD* mutant was impaired in gating molecular exchange upon H_2_O_2_ treatment (Figure S11). In addition to gating under certain stress conditions and providing a cargo-size cutoff, the complexity of the SJ architecture might also allow for the selection of specific cargo.

## Conclusion

In conclusion, our data suggest that cyanobacterial SJs are dynamic, gated cell-cell connections, which reversibly block intercellular molecular diffusion along the filament upon different types of stress (Figure 5). This challenges the concept of considering the cyanobacterial filament as a symplast, with SJs providing cytosolic continuity between the cells — analogous to plasmodesmata. SJs rather reveal striking similarities to metazoan gap junctions, because they are both gated by a dynamic conformational change of a proteinaceous macromolecular complex. Furthermore, just like SJs, gap junction closure is triggered by disruption of the membrane potential (Hervé and Derangeon, 2013; Obaid et al., 1983; Socolar and Politoff, 1971). Interestingly, the closure of gap junctions can be only partial (Ek-Vitorin and Burt, 2013), a phenomenon that might also exist in SJs, considering the ‘slow increase’ FRAP response (Figure 2b). Finally, gap junction and SJ closure might operate on a similar time scale, since we found that intercellular communication already ceased in less than four minutes after CCCP treatment of *Anabaena* cells (Figure S12).

**Figure 5:**
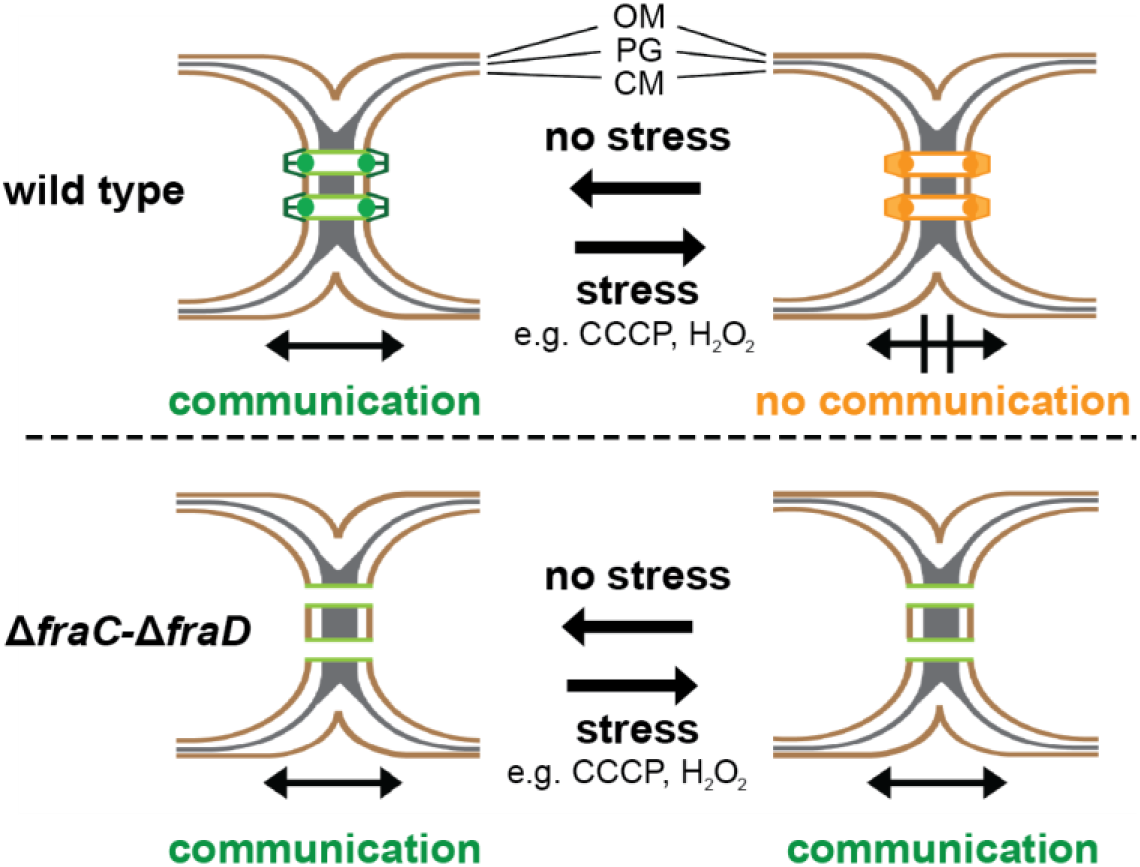
SJs reversibly gate cell-cell communication by a conformational change. SJs (green, open; orange, closed) of *Anabaena* are dynamic, gated cell-cell connections, which reversibly block intercellular molecular diffusion along the filament upon different types of stress. The Δ*fraD* and Δ*fraC-*Δ*fraD* mutants were missing the cap and plug modules, consistent with the inability to close SJs upon stress.

Cryotomography imaging of the cyanobacterial model organisms *Nostoc punctiforme* PCC 73102 and *Trichodesmium erythraeum* IMS101 showed similar SJ architectures (Figure S13). Together with the conservation of *fraC*, *fraD* and *amiC* genes in their genomes, this points towards a conserved SJ mechanism across diverse members of the phylum. The branching of the cyanobacterial order *Nostocales* (comprising the genus *Anabaena*) was estimated to date back more than two billion years ago (Schirrmeister et al., 2013). Our data thus provide a mechanistic framework for an ancient cell-cell connection structure, predating metazoan gap junctions by more than a billion years. The convergent evolution of a dynamic gated mechanism in such divergent lineages emphasizes the importance of controlling molecular exchange in order to stop communication under certain metabolic conditions, or upon predation or fragmentation. Avoiding leakage of the cytoplasm prevents the death of the entire multicellular organism.

## Acknowledgments

We thank P. Tittmann and C. Zaubitzer for technical support as well as ScopeM for instrument access at ETH Zürich. We thank Hannah Minas for help in preparing samples and analyzing the data. Jörn Piel and Anna Vagstaad are acknowledged for providing *Anabaena* and *Nostoc punctiforme* cultures for preliminary observations. We thank Ulrike Pfreundt and Roman Stocker for providing *Trichodesmium erythraeum* cultures. We thank Enrique Flores for providing FraC and FraC/FraD double mutants. Fabian Eisenstein is acknowledged for help with movies. GLW was supported by a Boehringer Ingelheim Fonds PhD fellowship. Work in Tübingen was supported by the Deutsche Forschungsgemeinschaft (SFB766). MP was supported by the Swiss National Science Foundation, the European Research Council and the Helmut Horten Foundation. Subtomogram averages were deposited in the Electron Microscopy Data Bank (accession numbers ##).

## Author contributions

GLW and AKK contributed equally. IM, KF and MP conceptualized the study. All authors designed experiments. GLW and AKK performed experiments. All authors analyzed data. All authors participated in writing the manuscript.

## Declaration of Interests

The authors declare no competing interests.

